## Supplementary Information for "EXCRETE enables deep proteomics of the microbial extracellular environment"

Files included in this document:
Supplementary Figures 1-6
Supplementary Tables 1-2
Separate SI excel sheets given for Supplementary Data 1-10

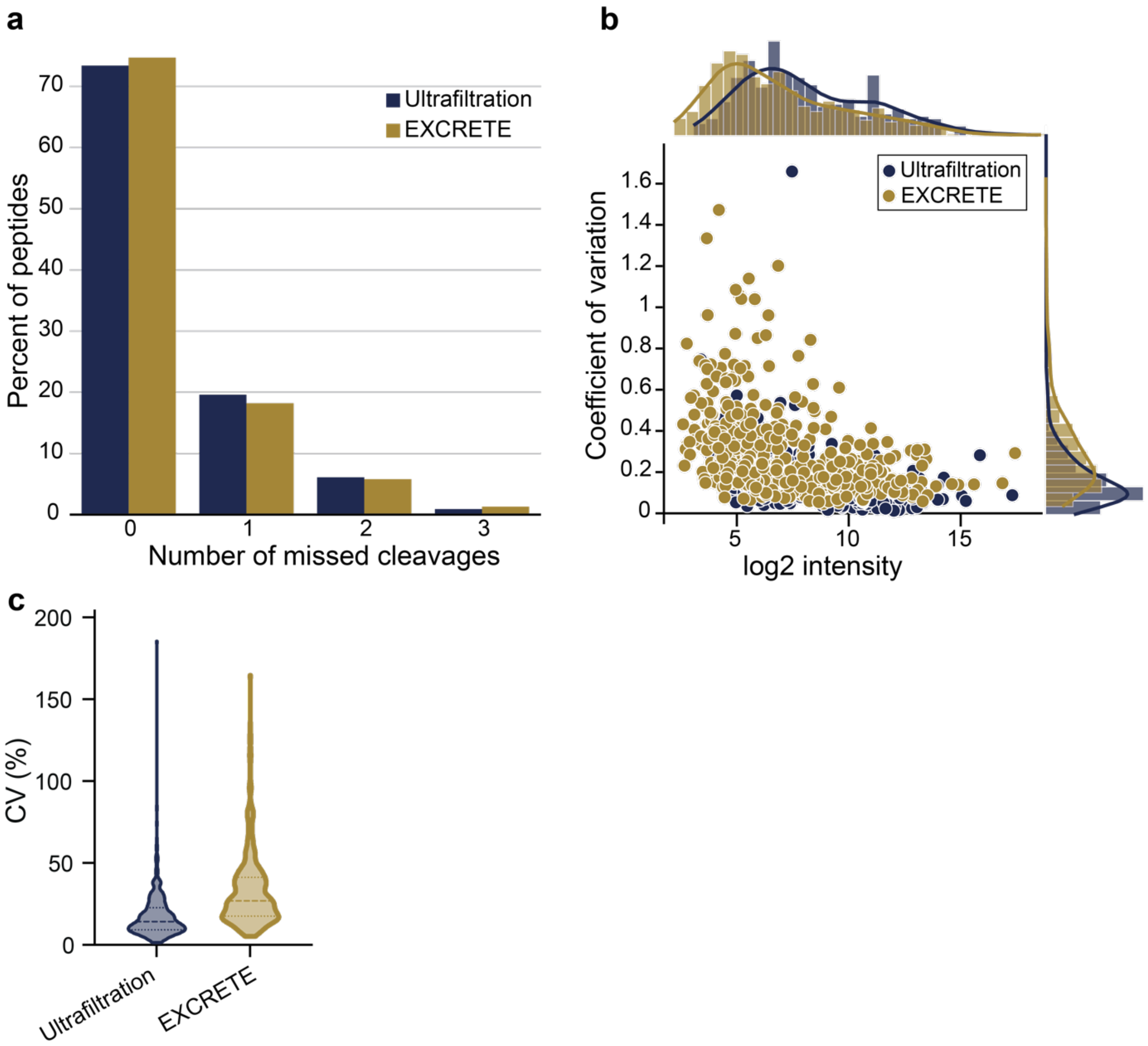


**Supplementary Figure 1** Benchmarking of EXCRETE against ultrafiltration-based exoproteomic sample preparation. **a** Percentage of peptides containing missed cleavages after digestion with Trypsin and Lys-c **b** Coefficients of variation (CVs) ordered by protein intensity. On the secondary x-axis histogram and density plots representing the frequency distribution of protein intensities are shown. On the secondary y-axis histogram and density plots representing the frequency distribution of CVs are shown. Dots represent means of biological replicates. **c** Coefficient of variation (CV) of proteins identified with ultrafiltration and EXCRETE.

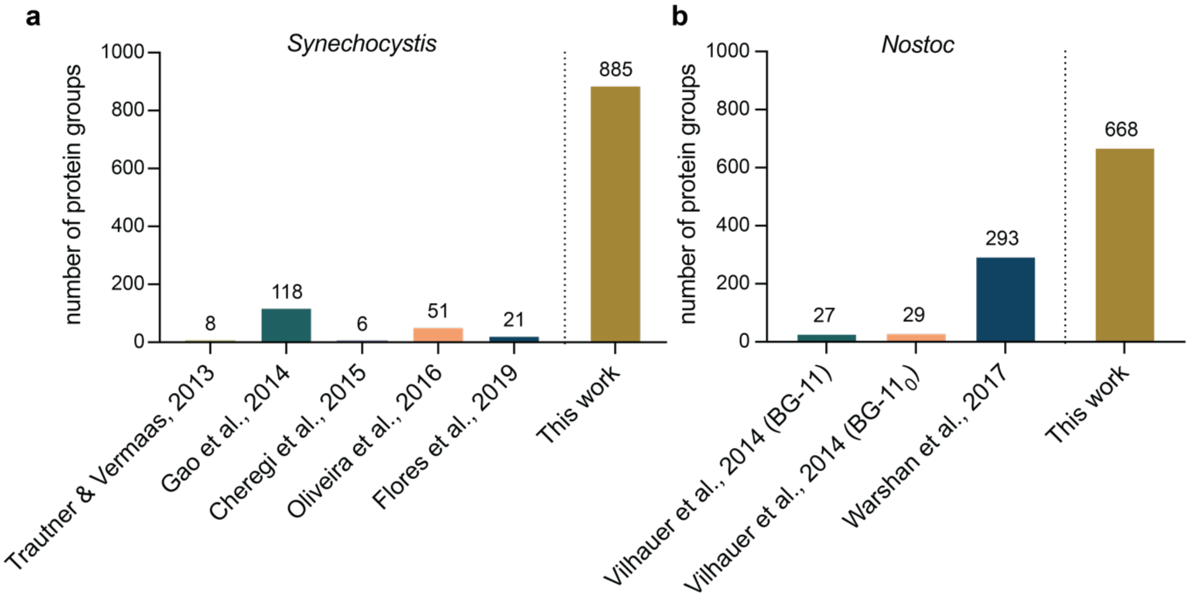


**Supplementary Figure 2** Comparison between the number of exoproteins identified in existing studies with the number identified in this study with EXCRETE.

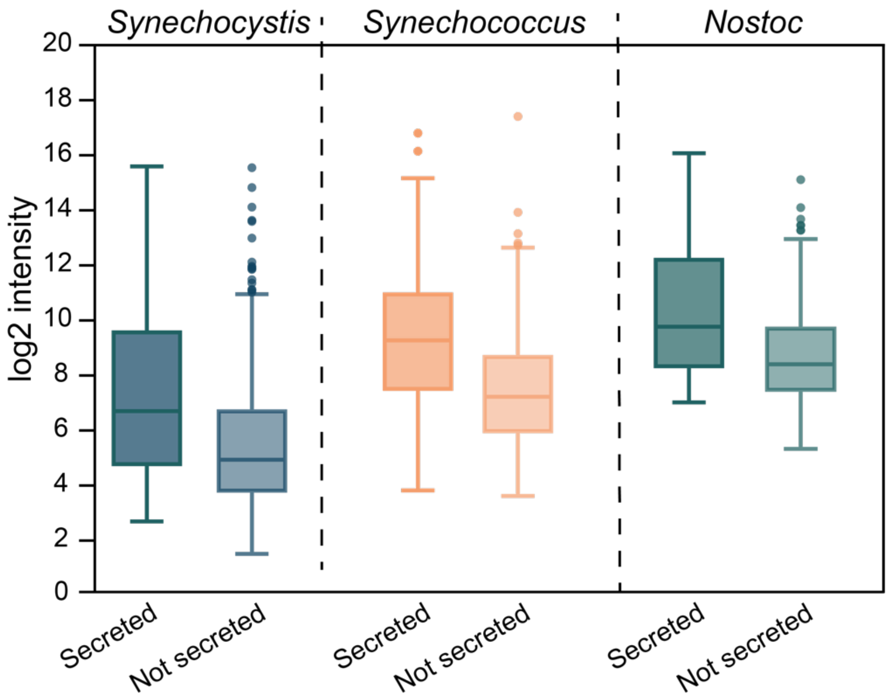


**Supplementary Figure 3** Total intensity of secreted and non-secreted proteins identified in *Synechocystis*, *Synechococcus* and *Nostoc*. Centre line of boxplots, median; box limits, upper and lower quartiles; whiskers, minimum to maximum values; dots, outliers.

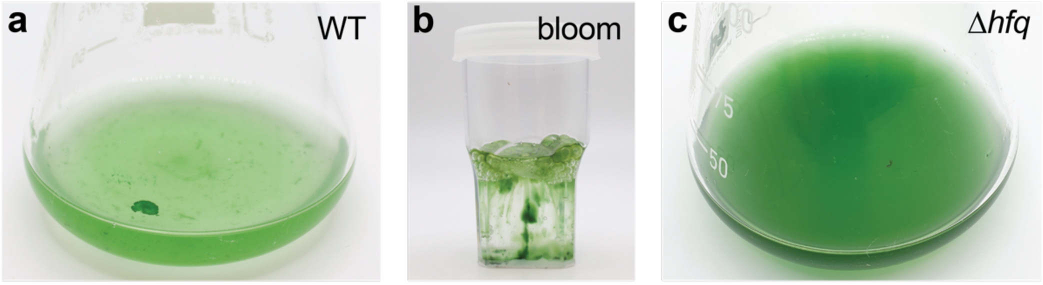


**Supplementary Figure 4** Representative pictures of *Synechocystis* cultures. **a** WT in standard conditions. **b** Bloom-like culture cultivated in high $CO_2$. **c** Δ*hfq* mutant.

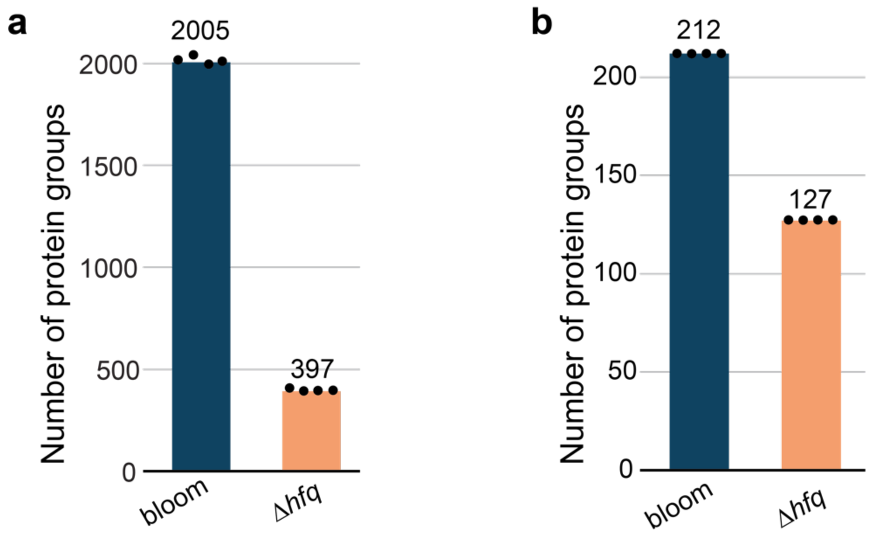


**Supplementary Figure 5** Number of total and secreted protein groups, after filtering and imputation, in the *Synechocystis* exoproteome in different conditions. **a, b** Number of total (**a**) and secreted (**b**) protein groups identified in a bloom-like culture and a Δ*hfq* mutant. Means are shown above the bars. Black dots represent biological replicates (n = 4).

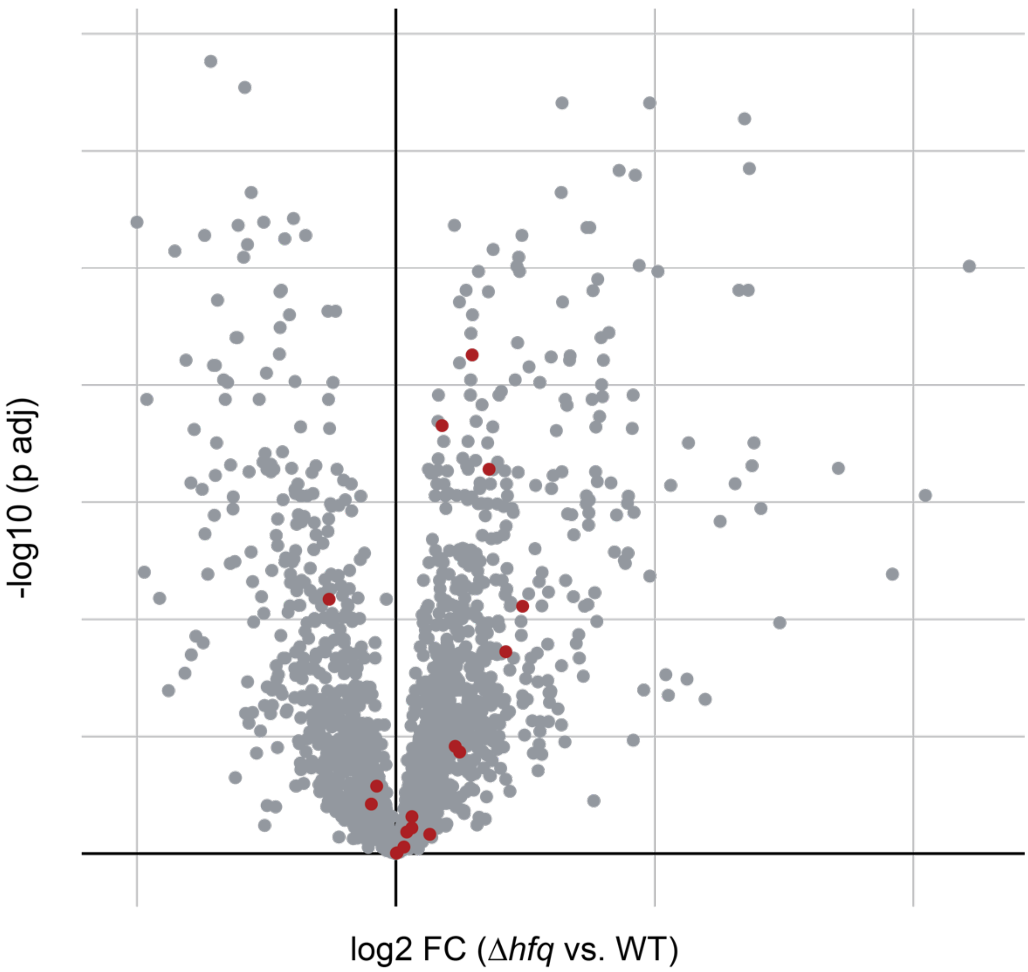


**Supplementary Figure 6** Volcano plot illustrating differential protein expression between proteins identified in the endoproteomes of WT *Synechocystis* and the Δ*hfq* mutant. Each dot on the plot represents an individual protein. Red dots indicate proteins that are absent from the *Synechocystis* secretome in the Δ*hfq* condition when compared to the WT conditions.

**Supplementary Table 1.** Putative T1SS substrates identified in the *Synechocystis* secretome.

| Protein ID (NCBI) | Protein ID (UniprotKB) | Protein name | Length (AA) | Locus tag | PosrtB localization | Signal peptide | PI | Gly (%) | Cys (%) |
|---|---|---|---|---|---|---|---|---|---|
| AGF52383.1 | NA | hypothetical protein | 4787 | NA | Extracellular | NA | 3.43 | 9.4 | 0.1 |
| AGF53432.1 | Q6ZEX5 | hypothetical protein | 3797 | slr5005 | Extracellular | NA | 4.41 | 11.7 | 0 |
| AGF52230.1 | P74440 | integrin alpha subunit domain-like protein | 4199 | slr0408 | Extracellular | NA | 3.99 | 10.9 | 0 |
| AGF52739.1 | Q55365 | endo-1,4-beta-glucanase | 1070 | slr0897 | Extracellular | NA | 4.6 | 13.7 | 0 |
| AGF53312.1 | P74647 | hypothetical protein | 1771 | sll0723 | Extracellular | NA | 4.37 | 10.9 | 0 |
| AGF53314.1 | P74649 | leukotoxin LtA | 1290 | sll0721 | Extracellular | NA | 4.03 | 12.8 | 0 |
| AGF50742.1 | P73032 | hypothetical protein | 1749 | slr1753 | Extracellular | Sec lipo | 3.81 | 10.2 | 0.2 |
| AGF50645.1 | P72939 | alkaline phosphatase | 1409 | sll0654 | Periplasm | NA | 3.93 | 9.9 | 0 |
| AGF51323.1 | P73590 | integrin alpha- and beta4-subunit domain-like protein | 3016 | slr1403 | Extracellular | NA | 3.98 | 14.7 | 0 |
| AGF53129.1 | Q55489 | hypothetical protein | 948 | sll0499 | Outer membrane | NA | 4.6 | 6.9 | 0.1 |
| AGF50804.1 | P73089 | fat protein | 1965 | slr2046 | Extracellular | NA | 3.61 | 7.8 | 0 |

**Supplementary Table 2.** COG classification of the 54 proteins upregulated in the *Synechocystis* secretome in the Δ*hfq* condition in comparison to the WT condition.

| **COG category** | **%** |
|---|---|
| Not attributed | 30 |
| Cell envelope biogenesis | 19 |
| Function unknown | 13 |
| Carbohydrate transport and metabolism | 9 |
| Secondary metabolism | 9 |
| Signal transduction mechanism | 9 |
| Trafficking, secretion, and vesicular transport | 6 |
| Amino acid transport and metabolism | 4 |
| Cell cycle control and cell division | 4 |
| Transcription | 2 |
| Replication, recombination and repair | 2 |
| Energy production and conversion | 2 |
| Defense mechanisms | 2 |
| Inorganic ion transport and metabolism | 2 |
